# Seedless fruit in *Annona squamosa* L. is monogenic and conferred by *INO* locus deletion in multiple accessions

**DOI:** 10.1101/2022.10.25.513714

**Authors:** Bruno Rafael Alves Rodrigues, Charles S. Gasser, Samy Pimenta, Marlon Cristian Toledo Pereira, Silvia Nietsche

## Abstract

Understanding the genetic basis and inheritance of a trait facilitates the planning of breeding and development programs of new cultivars. In the sugar apple tree (*Annona squamosa* L.), the mechanism of the desirable seedless trait in the Thai seedless (Ts) and Brazilian seedless (Bs) accessions was associated with a deletion of the *INNER NO OUTER* (*INO*) locus. Genetic analysis of F_1_, F_2_ and backcross descendants of crosses of Bs to fertile wild-type varieties showed that seedlessness was recessive and monogenic. Whole genome sequencing of a third accession, Hawaiian seedless (Hs), identified a 16 kilobase deletion including *INO*. The finding of an identical deletion in Ts and Bs indicated a common origin among genotypes, from a single deletion event. Analysis of microsatellite markers could not preclude the possibility that all three accessions are vegetatively propagated clones. The sequence of the deletion site enabled formulation of a codominant assay for the wild-type and mutant genes that validated the *INO* gene deletion as the cause of seedless trait, and can be used in the selection of new seedless varieties. The study findings and obtained progenies should be useful in breeding and introgression programs of the trait into elite sugar apple lines and into other *Annonas* by means of interspecific crossings.

## Introduction

The global fruit market has grown appreciably in the past decade. The absence of seeds from fruit that is consumed has become one of the most appreciated traits by consumers. Reducing seed content without changing the size of the fruit is one of the main objectives of the cultivation of many fruit trees. The goal is to enhance the experience of fruit consumption by consumers and improve the quality of fruits for food processing (Varoquaux et al., 2000).

Sugar apple (*Annona squamosa* L.) (2n = 14) is a tropical fruit species in the family Annonaceae. Seedless fruits have been described for some spontaneous mutants of *A. squamosa* whose origin is still unknown (Pinto et al., 2005; Wongs-Aree and Noichinda, 2011; Lora et al., 2018; Pereira and Borém, 2021). They include the Cuban cultivar Cuban seedless (Cs), with good fruit characteristics but lower productivity than fertile cultivars (Araújo et al., 1999); Brazilian seedless (Bs) originally identified in northeast Brazil, which produces small, asymmetric fruits that frequently perish (Santos, 2014); the Thai seedless (Ts) mutant that produces normal size fruits (Lora et al. 2011) among other types with apparently similar fruits, such as in the Philippines and Hawaii (Pinto et al., 2005; this work).

Lora et al. (2011) were among the first researchers to examine details of the seedless trait in *A. squamosa* in Ts. This variety produces fruit following pollination and fertilization. The authors demonstrated that seedlessness results from a defect in ovule development where the outer of the two integuments sheathing the nucellus fails to form. This defect directly mirrors the effect of *INNER NO OUTER* (*INO*) loss of function mutants in *Arabidopsis thaliana* (Villanueva et al., 1999). *INO* encodes a putative transcription factor belonging to the *YABBY* family. Members of this family are involved in the determination of abaxial identity in a variety of plant organs (Sawa et al., 1999; Siegfried et al., 1999; Bowman, 2000). Lora et al. (2011) isolated an *A*. *squamosa INO* ortholog and demonstrated the association of the Ts mutant with an apparent deletion of the *INO* gene, indicating a candidate gene for the seedless trait.

The Bs variety was also evaluated in a breeding program undertaken at the State University of Montes Claros. Results showed that the absence of seeds was also associated with failure in the development of the external integument of the ovule and in the development of seeds, similar to that observed in Ts (Mendes et al., 2012; Santos et al., 2014). In addition, preliminary studies of inheritance of the stenospermocarpic absence of seeds involving F1 progenies in Bs (*A. squamosa* wild type x Bs) also indicated the same expected recessive nature for the mutation as in Ts (Nassau et al., 2021). However, it was not known whether the molecular basis of the *INO* deletion described is widespread in *A. squamosa* and may be responsible for the case of aspermia described in Bs.

The present study addressed the following questions: (i) Is the inheritance of the presence/absence of seeds in *A. squamosa* monogenic?; (ii) What are the molecular details of the *INO* gene deletion?; (iii) Is there a complete correspondence of homozygosity for the deletion with the seedless phenotype; (iv) What is the relationship between the known seedless accessions, and (v) Can codominant molecular markers specific to *INO* be designed for use in assisted selection?

## Materials and Methods

### Plant material

Wild-types (fertile) M_1_, M_2_ and M_3_ and seedless Bs lines were previously described by de Souza et al. (2010), and Ts was described by Lora et al. (2011). For whole genome sequencing a wild-type sugar apple fruit was purchased from a retail source in the United States, seeds from the fruit were planted, and one plant grown in the UC Davis Conservatory was sampled for sequencing with voucher herbarium samples stored as DAV225058 and DAV225059. The Hawaiian seedless (Hs) line was obtained as budwood from Frankie’s Nursery (Waimanalo, HI), grafted to a wild-type *A. squamosa* rootstock and grown in the UC Davis Conservatory with a voucher sample stored as DAV225060.

For genetic inheritance studies, plants (M_1_, M_2_, M_3_ and Bs) were grown at the Experimental Farm and molecular analysis were performed at Molecular Biology laboratory of the State University of Montes Claros, latitude 15°48’09’’S, longitude 43°18’32’’W and altitude 516 m. For phenotypic characterization of seedless versus seeded (fertile) two strategies were applied: (a) fruits were harvested, pulped, and examined for the presence or absence of seeds (Supplemental Figure S1); or (b) flowers either fresh or fixed in formalin acetic acid-alcohol (FAA) were dissected to separate the ovules from the carpel tissue. Isolated ovules were observed with a stereomicroscope (Nikon, SMZ800) and photographed by a digital camera (Sony, Dsc-W350). The wild-type ovules present a domed shape opposite the funiculs, while the mutant ovules come to a point at this position (Supplemental Figure S2).

Filial generations (F_1_), self-fertilization (F_2_), backcrosses with wild-types parents M_1_, M_2_, M_3_ (BC_M_), and backcrosses with mutant parent Bs (BC_Bs_) were obtained. Segregations were evaluated for conformity to predicted ratios with the Chi-square test (χ^2^) using the Genes statistical software (Cruz, 2016).

### Polymerase chain reaction (PCR) analysis

DNA was extracted from young leaf samples with hexadecyltrimethylammonium bromide buffer (CTAB) as described by Doyle and Doyle (1990) and separated from polysaccharides as described by Cheung et al., (1993). Primers used in PCR are listed in Supplemental Table S1. PCR was performed with DreamTaq (ThermoFisher, Waltham, MA) and the included reagents with an initial denaturation at 94°C for 3 min; 35 cycles with denaturation at 94°C for 30 sec, annealing at 56°C for 30 sec, and extension at 72°C for 1.5 min; and a final extension of 72°C for 4 min. For reactions using the AsINODel primers a 60° annealing temperature was used.

PCR products were electrophoresed on 1.2 % (w/v) agarose buffered with 1× TBE (89 mM Tris-base, 89 mM boric acid, 2 mM EDTA, pH 8.0) or SB (10 mM Sodium Borate pH 8.0) and DNA visualized by staining with ethidium bromide (2 μg/ml) an illumination with ultraviolet light.

For sequencing, PCR products were processed with ExoSAP-IT (ThermoFisher, Waltham, MA) or Quiapure (Quiagen, Germantown, MD) and sequenced using amplification primers on an ABI 3500 or 3730 genetic analyzer (ThermoFisher, Waltham, MA) at Análises Moleculares Ltda. (Centro de Biotecnologia, UFRGS, Porto Alegre, RS) or the University of California Davis CBS DNA Sequencing Facility (Davis, CA).

### Whole genome shotgun sequencing

DNA for whole genome sequencing was isolated from young leaves by grinding in 100 mM TRIS-Cl, 20 mM EDTA, 1.4 M NaCl, 2% (w/v) CTAB, 1% each polyvinylpyrrolidone and sodium metabisulfite pH 8.0. Samples were treated with 70 μg/ml RNAaseA (ThermoFisher, Waltham, MA), extracted with 1:24 (v/v) mixture of isoamyl alcohol and chloroform, and precipitated with isopropanol. Samples were dissolved in 10 mM TRIS pH 8.0, 1 mM EDTA, adjusted to 0.3 M Na Acetate, pH 4.8, precipitated with 2 volumes of ethanol and dissolved in 10 mM TRIS pH 8. Wild-type *A. squamosa* DNA was processed and sequenced at the University of California, Davis Genome Center (Davis, CA). For PacBio (Menlo Park, CA) sequencing, DNA fragments greater than 10 kb were selected by BluePippin (Sage Sciences, Beverly, MA) electrophoresis and were sequenced on a PacBio RSII or Sequel Single Molecule, Real-time device. This resulted in 2.46 million reads with an average read length of 8 kb comprising more than 29 Gbases, or approximately 37 X genome representation. For Illumina (San Diego, CA) sequencing the DNA was sheared and fragments of an average size of 400 bp were selected and sequenced on a HiSeq 4000 apparatus by the paired-end 150 bp method (PE150) resulting in approximately 390 million sequences. The sequences were trimmed of poor quality regions and primer sequences with Sickle (Joshi and Foss, 2011) and Scythe (Buffalo, 2011), respectively resulting in 229 Gbases or approximately 124 X genome representation. Hs DNA was similarly processed and sequenced by QuickBiology (Pasadena, CA) resulting in 408 million sequences and approximately 130 X genome representation.

### Genome sequence assembly and analysis

The PacBio reads of wild-type DNA were assembled using Canu (Koren et al., 2017) with default settings, producing 3519 contigs. The wild-type Illumina reads were aligned with the assembly using BWA MEM (Li, 2013) with default settings, and Pilon (Walker et al., 2014) used the alignment to correct the contigs, changing 148k single nucleotide errors and adding a net of more than 1.8 Mbases of insertions for a final assembly of 707.7 Mbases with an average contig length of 201 kb and an N90 of 93.9 kb. A BLAST (Altschul et al., 1990) search with the known *A. squamosa INO* gene sequence (Lora et al., 2011) identified a 587 kb contig (tig00001115) containing the *INO* gene (Supplemental Figure S3). BWA MEM aligned the Hs Illumina reads with the assembly and Tablet (Milne et al., 2013) was used to examine the alignment with the 587 kb contig containing INO. BLAST was used to search one half of the set of Hs Illumina sequence reads for those extending across a detected deletion and the resulting sequences were aligned and assembled using Sequencher 5.4.1 (Gene Codes, Ann Arbor, MI).

### Genetic diversity using SSR markers

Genetic diversity was assessed among varieties of seedless sugar apple Bs, Ts, and Hs, with the fertile parent M_2_ as a contrasting control. Sixty-seven pairs of SSR microsatellite markers, described for *A. cherimola* were used, with fifteen having been described by Escribano et al. (2004) and fifty-two by Escribano et al. (2008).

DNA extraction was performed as described above for the markers association of seedless trait with INO deletion. Amplification utilized an initial denaturation at 94°C for 1 min; 35 cycles at 94°C denaturation for 30 sec, annealing at 48 to 57°C depending on the primer; and extension at 72°C for 1 min; and a final extension of 72°C for 7 min. The amplification products were separated by 3.0% agarose gel electrophoresis buffered, stained and visualized as above.

To calculate diversity, the amplification data of the SSR primers were converted into numerical code per locus for each allele. The presence of a band was designated by 1 and the absence by 0. Although the microsatellite markers can be codominant, molecular analyses of the locus were performed based on the presence/absence of each amplified fragment. The established binary matrix was used to obtain estimates of genetic similarities between genotype pairs, based on the Jaccard coefficient. The Genes statistical program (Cruz, 2016) was used for data processing.

### Procedures adopted chronologically

The present study was developed according the following chronological events: (1) obtaining the segregating generations (F_2_, BC_M_ and BC_Bs_) from the crossing of the fertile parents M_1_, M_2_ and M_3_ with the mutant Bs (2018-2019); (2) application of the dominant marker LMINO primer-set in order to associate the seedless trait with *INO* deletion in the segregating generations (F_1_, F_2_, BC_M_ and BC_Bs_) (2019-2020); (3) sequencing and identification of the *INO* gene deletion region and design the codominant primer AsINODel (2021); (4) AsINODel marker validation and phenotypic characterization of plants (F_2_, BC_M_ and BC_Bs_) in the field conditions (2021-2022).

## Results

### Seedless trait inheritance

The results of the phenotypic analysis of the parents M_2_ and Bs, F_1_, F_2_, backcrosses with the wild-type parent M_2_ (BC_M_) and with mutant parent Bs (BC_Bs_) are displayed in Table 1. In generation F_1_, all individuals presented fruits with seeds. In the F_2_ population, among the plants in reproductive stage during the evaluation period, 48 formed fruits with fully developed seeds and ten presented only seed rudiments, characterized by the absence of seeds (Supplemental Figure S1). Considering segregation hypotheses expected for one, two and three genes (Supplemental Table S2), the Chi-square test revealed that the trait under study segregated at a 3:1 ratio (presence/absence), consistent with a monogenic inheritance. These results were corroborated by data from backcrosses with the parent M_2_ (BC_M_), where all plants that produced fruits had seeds, consistent with the 1:0 ratio, while the plants evaluated through backcrossing with the mutant parent Bs (BC_Bs_), showed segregation of 1:1 for presence and absence of seeds. Taken together, these results corroborate the monogenic inheritance found in the analyses of F_2_ generations, indicating that a single recessive locus controls the seedless trait in Bs *A. squamosa*.

**Table 1.** Phenotypic analysis of the segregation of the seed presence/absence in fruits of F_1_ and F_2_ generations, BC_M_ and BC_Bs_ backcrosses of the M_2_ family in *A. squamosa.* Janaúba-MG, Brazil.

| Parents and Generations | Observed proportion | | Expected ratio | $\chi^2$ | *p*-value |
| --- | --- | --- | --- | --- | --- |
|  | Presence | Absence |  |  |  |
| Parents |  |  |  |  |  |
| M<sub>2</sub> | 1 |  |  |  |  |
| Bs |  | 1 |  |  |  |
| Generation F<sub>1</sub> |  |  |  |  |  |
| M<sub>2</sub> × Bs | 17 | 0 | 1:0 |  |  |
| Generation F<sub>2</sub> |  |  |  |  |  |
| F<sub>1</sub>(M<sub>2</sub> × Bs) × F<sub>1</sub>(M<sub>2</sub> × Bs) | 48 | 10 | 3:1 | 1.86 | 0.172<sup>ns</sup> |
| Backcrossing |  |  |  |  |  |
| BC<sub>M</sub> |  |  |  |  |  |
| F<sub>1</sub>(M<sub>2</sub> × Bs) × M<sub>2</sub> | 12 | 0 | 1:0 |  |  |
| BC<sub>Bs</sub> |  |  |  |  |  |
| F<sub>1</sub>(M<sub>2</sub> × Bs) × Bs | 8 | 6 | 1:1 | 0.286 | 0.592<sup>ns</sup> |

<sup>ns</sup> Non-significant values at the 5% significance level, $\chi^2$ Chi-square test estimate.

### Association of seedless trait with *INO* deletion

Previously described molecular markers for the presence of the *INO* gene (Lora et al., 2011) were tested on parents M_1_, M_2_, M_3_, and Bs and displayed the expected band patterns. These markers generated amplification products only in the three wild-type parents, with no amplification of any fragment in Bs for any of the primer pairs used (Supplemental Figure S4A).

The dominant marker LMINO primer-set were also used to amplify DNA from F_1_ plants obtained from crosses between genotypes of *A*. *squamosa* (M_1_, M_2_, and M_3_) with the mutant Bs (Table 1). All evaluated F_1_ individuals produced fruits with seeds in the field and amplified the products with all primer pairs, as shown in the Supplemental Figure S4B.

The same procedure was applied in order to genotype segregating generations (F_2_, BC_M_ and BC_Bs_) in seedling stage in the nursery (Table 2). Figure 1 shows a sample of individuals amplified with the LMINO1/2 primers and the results confirm the discriminatory capacity of those genetic markers (Figure 1A, B and C). The field confirmation of presence/ absence of seeds in the fruits in these generations F_2_, BC_M_ and BC_Bs_ were obtained later (Table 1).

**Figure 1.**
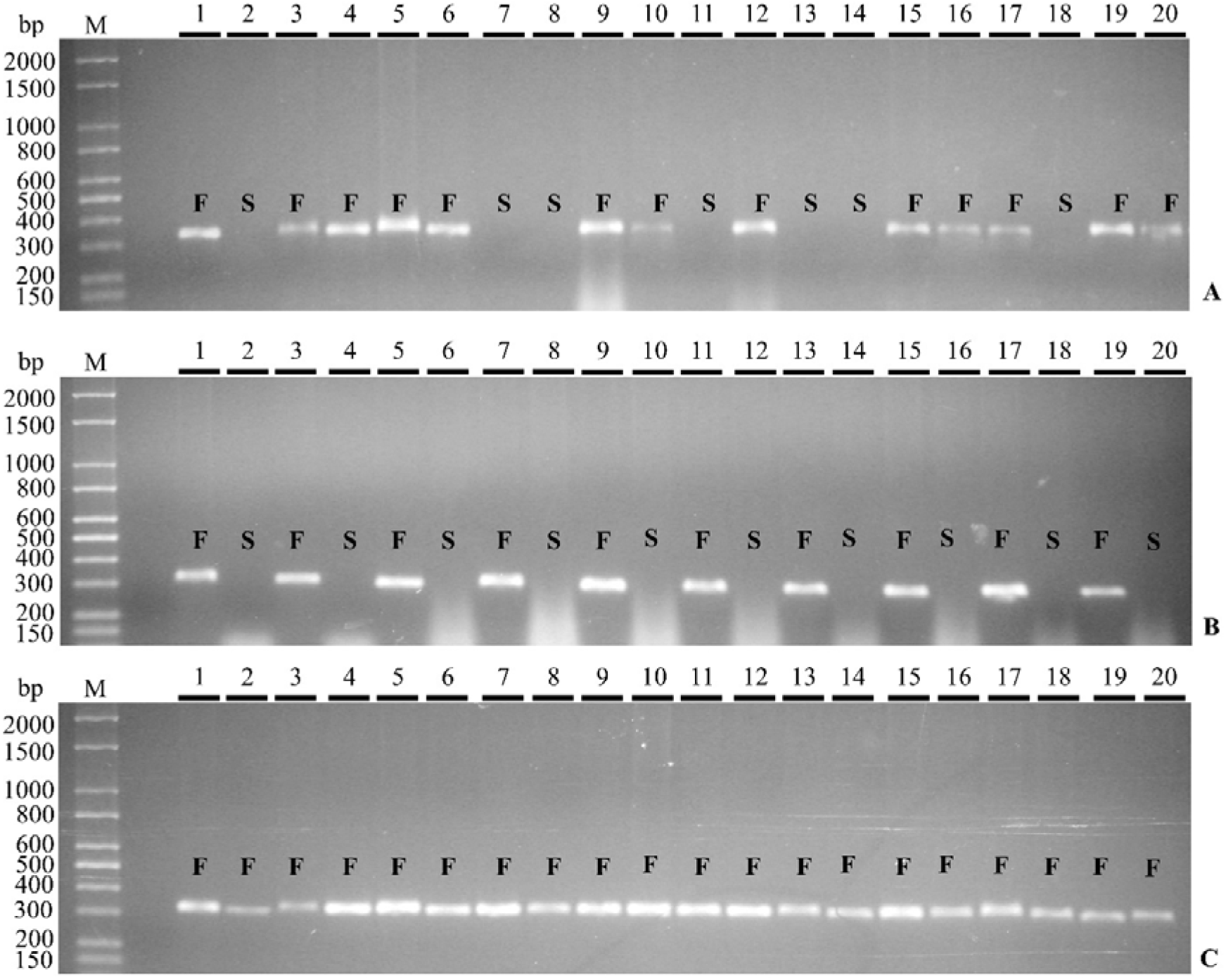
Amplification products for primer pair LMINO1/2 obtained from DNA samples from segregating generations: A) F_2_, B) BC_Bs_ C) BC_M_ in 1.2% agarose gels in 1× TBE buffer. M bp: molecular weight marker in base pairs. Lanes 1 to 20 contain the respective amplified fragments from the 20 genotypes. S: seedless; F: fertile (S and F are phenotypic determinations).

**Table 2.** Molecular analysis of segregation of dominant primers in parents (M_1_, M_2_, M_3_ and Bs), generations F_1_, F_2_, BC_M_ and BC_Bs_ in *A. squamosa,* Janaúba-MG, Brazil.

| Parents and Generations | Observed proportion | | Expected ratio | $\chi^2$ | *p*-value |
| --- | --- | --- | --- | --- | --- |
|  | Presence | Absence |  |  |  |
| Parents |  |  |  |  |  |
| M<sub>1</sub> | 1 |  |  |  |  |
| M<sub>2</sub> | 1 |  |  |  |  |
| M<sub>3</sub> | 1 |  |  |  |  |
| Bs |  | 1 |  |  |  |
| Generation F<sub>1</sub> |  |  |  |  |  |
| M<sub>1</sub> × Bs | 9 | 0 | 1:0 |  |  |
| M<sub>2</sub> × Bs | 17 | 0 | 1:0 |  |  |
| M<sub>3</sub> × Bs | 6 | 0 | 1:0 |  |  |
| Generation F<sub>2</sub> |  |  |  |  |  |
| F<sub>1</sub>(M<sub>1</sub> × Bs) × F<sub>1</sub>(M<sub>1</sub> × Bs) | 99 | 43 | 3:1 | 2.113 | 0.146<sup>ns</sup> |
| F<sub>1</sub>(M<sub>2</sub> × Bs) × F<sub>1</sub>(M<sub>2</sub> × Bs) | 190 | 71 | 3:1 | 0.676 | 0.411<sup>ns</sup> |
| F<sub>1</sub>(M<sub>3</sub> × Bs) × F<sub>1</sub>(M<sub>3</sub> × Bs) | 69 | 33 | 3:1 | 2.941 | 0.086<sup>ns</sup> |
| Heterogeneity |  |  |  | 1.182 | 0.553<sup>ns</sup> |
| Backcrosses |  |  |  |  |  |
| BC<sub>M</sub> |  |  |  |  |  |
| F<sub>1</sub>(M<sub>1</sub> × Bs) × M<sub>1</sub> | 63 | 0 | 1:0 |  |  |
| F<sub>1</sub>(M<sub>2</sub> × Bs) × M<sub>2</sub> | 67 | 0 | 1:0 |  |  |
| F<sub>1</sub>(M<sub>3</sub> × Bs) × M<sub>3</sub> | 61 | 0 | 1:0 |  |  |
| BC<sub>Bs</sub> |  |  |  |  |  |
| F<sub>1</sub>(M<sub>1</sub> × Bs) × Bs | 48 | 61 | 1:1 | 1.550 | 0.213<sup>ns</sup> |
| F<sub>1</sub>(M<sub>2</sub> × Bs) × Bs | 57 | 59 | 1:1 | 0.034 | 0.852<sup>ns</sup> |
| F<sub>1</sub>(M<sub>3</sub> × Bs) × Bs | 26 | 30 | 1:1 | 0.286 | 0.593<sup>ns</sup> |
| Heterogeneity |  |  |  | 0.586 | 0.746<sup>ns</sup> |

<sup>ns</sup> Non-significant values at the 5% significance level, $\chi^2$ Chi-square test estimate.

In the F_2_ generations of the three crosses (M_1_ × Bs, M_2_ × Bs and M_3_ × Bs), there was a segregation of the products of the amplification of the LMINO markers that correlated exactly with the presence/absence of seeds. Fertile plants in this generation uniformly produced an amplification product with the LMINO1/2 primer set, while plants producing no product (indicating homozygosity for the *INO* gene deletion) produced only seedless fruit (Figure 1A). The same complete cosegregation pattern seen in F_2_ individuals for the presence/absence of seeds and PCR product was also observed in backcross populations of BC_Bs_ (Figure 1B). For BC_M_ backcross plants, the formation of *INO* amplification products was observed in all DNA samples tested (Figure 1C) for these uniformly fertile/seed bearing plants.

The χ^2^ test was performed with the data generated in the molecular analysis to confirm the segregation of the dominant amplification (Table 2). F_1_ plants displayed the expected genotypic ratio of 1:0 (presence/absence of amplified band) that had been linked to the trait of seeded fruits. In F_2_ generations, six segregation hypotheses expected for one, two and three genes were tested (Supplemental Table S3). Considering a significance of 5% probability, the frequencies of genotypes fit a ratio of 3:1, but allowed rejection of the other predicted ratios, confirming the hypothesis that a single locus confers the phenotype for the trait under study, with the dominant allele responsible for the presence and the recessive allele for the absence of the amplification product.

To identify the homogeneity between the F_2_ crossings (M_1_ × Bs, M_2_ × Bs, and M_3_ × Bs), statistical techniques were applied to verify whether the differences observed in the results could be explained by chance or not. The heterogeneity test was not significant and indicated, with a 55% likelihood, that the results of the χ^2^ were consistent for the populations of the three families studied, confirming the expected segregation (Table 2).

To further support the hypothesis of segregation in F_2_ generations, BC_M_ and BC_Bs_ backcrosses were used. Similarly, the heterogeneity of segregation between the families of the BC_Bs_ backcrossing was not significant (Table 2). BC_M_ and BC_Bs_ progenies analysed separately displayed segregation in a manner consistent with the hypothesis of a single gene. In BC_Bs_ backcrossing, carried out between generations F_1_ and the parent Bs, the proportion was close to 1:1 presence/absence of seeds in the fruits. χ^2^ test was applied and the deviations between the observed and expected frequencies were not significant. In the BC_M_ backcrossing between generations F_1_ and parents (M_1_, M_2_, and M_3_), the proportion was 1:0 presence/absence of seeds in the fruits. These results confirmed the monogenic inheritance found in the analyses of F_2_ generations consistent with a single recessive allele being responsible for the seedless trait in *A. squamosa* considering the 3:1 segregation hypothesis.

### The nature of the *INO* gene deletion event

Whole genome shotgun sequencing was used to determine the characteristics of the *INO* gene deletion event. A draft wild-type *A. squamosa* genome was assembled through sequencing of total DNA isolated from a plant grown from seed derived from commercially available *A. squamosa* fruit. Genomic DNA was sequenced by both long-read (PacBio (Menlo Park, CA) Single Molecule, Real-Time sequencing) and short read (Illumina (San Diego, CA)) paired-end 150 base (PE150) methods. The long reads were assembled into a draft sequence that was corrected with the higher coverage short read sequences. The resulting assembly comprised 707 Mbases of DNA in 3,519 contigs, with average contig length of 201 kb. A BLAST (Altschul et al., 1990) search with a previously published *A. squamosa INO* gene sequence (GenBank GU828033.1) was used to identify a 587 kb contig that included the *INO* gene (Supplemental Figure S3). Total Hs *A. squamosa* DNA was used to produce a second short-read sequence set (~140-fold redundant) and this was aligned with the assembled wild-type sequence. Visualization of the alignment of the Hs sequences with the 587 kb contig including *INO* revealed a clear absence of reads over a region of 16,020 bp indicating a 16 kb deletion that included the *INO* gene (Figure 2A). The alignment program truncates read sequences where they do not align with the reference sequence, so a deletion or a deletion with a heterologous insertion would appear similar in this visualization. Truncated sequence reads at the immediate borders of the putative deletion region were used to perform a BLAST search of the Hs sequence reads to identify untruncated read sequences. Identified sequence reads from the two sides of the deletion region overlapped allowing us to assemble the Hs sequence in this region (Figure 3). This showed that the lesion was in fact a clean deletion of the region deficient in read sequences with no inserted DNA and no sequence duplication at the deletion borders (Figure 2B). The deleted region included the entire *INO* gene and part of the first coding region of another putative gene encoding a possible UDP-N-acetylglucosamine-N-acetylmuramyl-pyrophosphoryl-undecaprenol N-acetylglucosamine protein. The genomic region containing *INO* in wild-type and the corresponding region from Hs were deposited in GenBank as accessions ON248606 and ON248607, respectively.

**Figure 2.**
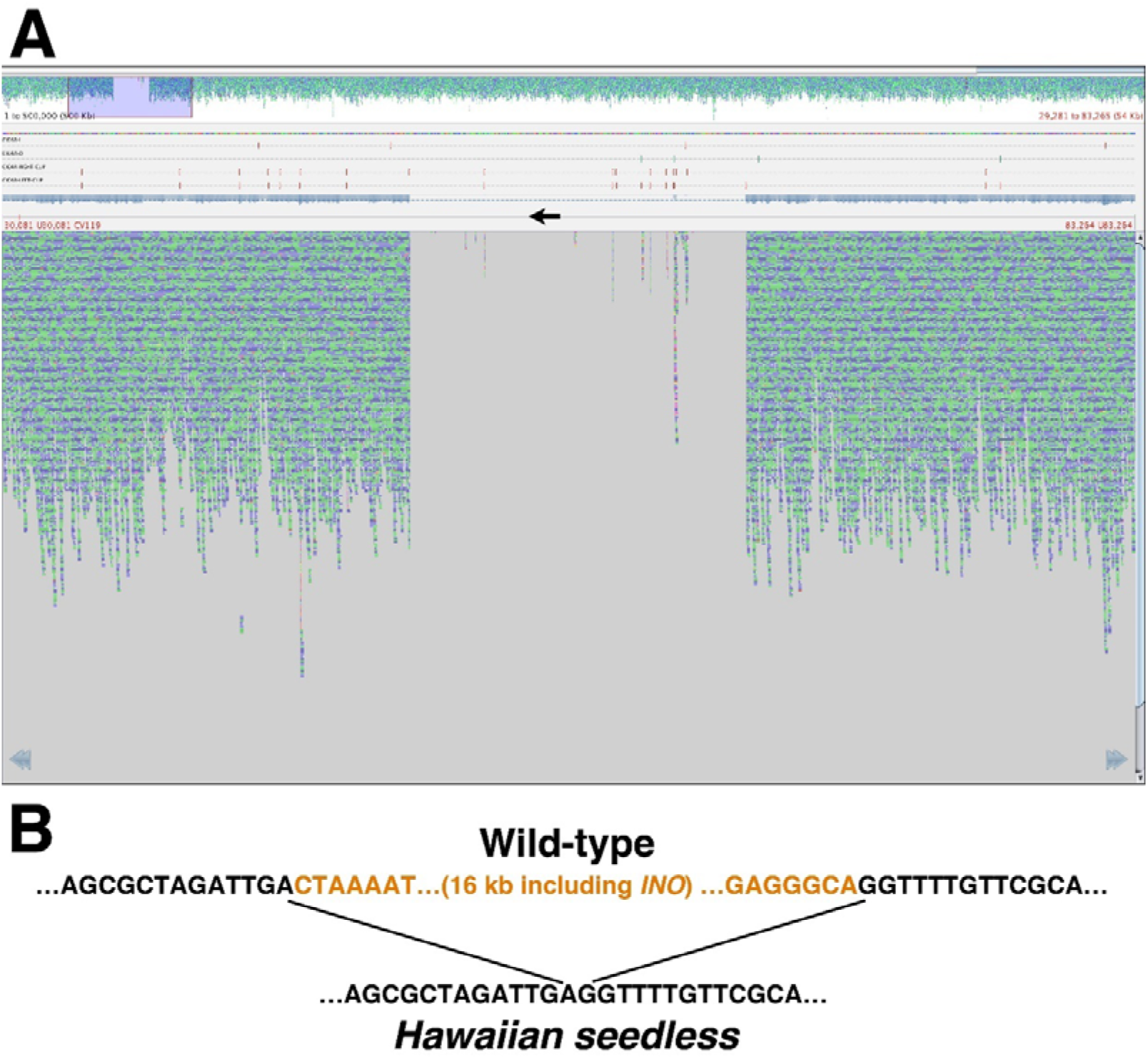
Identification of the *INO* deletion region. A) Screen capture of Tablet (Milne et al., 2013) visualization of the alignment of Hs Illumina sequence reads with the 587 kb contig of *A. squamosa* wild-type sequence that contains the *INO* gene. At top is a view of 500 kb of the alignment with each of the aqua or blue colored dots representing one 150 base read of the Hs sequence. A region including a deficiency of aligning reads is visible and is highlighted by a translucent purple box. The lower portion of the panel is an expanded view of the highlighted region where each small colored line again depicts a 150 base Hs read aligning with the wild-type reference sequence. The near complete absence of aligning reads in the center of this region (barring a small number of reads aligning with repeated sequences within this region) shows a clear deletion in the Hs sequence relative to the wild-type reference sequence. The black arrow edited onto the figure depicts the location of the *INO* gene transcribed region in the wild-type reference sequence. B) Depiction of the wild-type sequence compared to the sequence of the corresponding region in Hs as determined by assembling Hs Illumina reads that overlapped the deletion junction. A deletion of 16,020 bp was observed with no duplication of sequence at the junction site and no additional inserted sequence.

**Figure 3.**
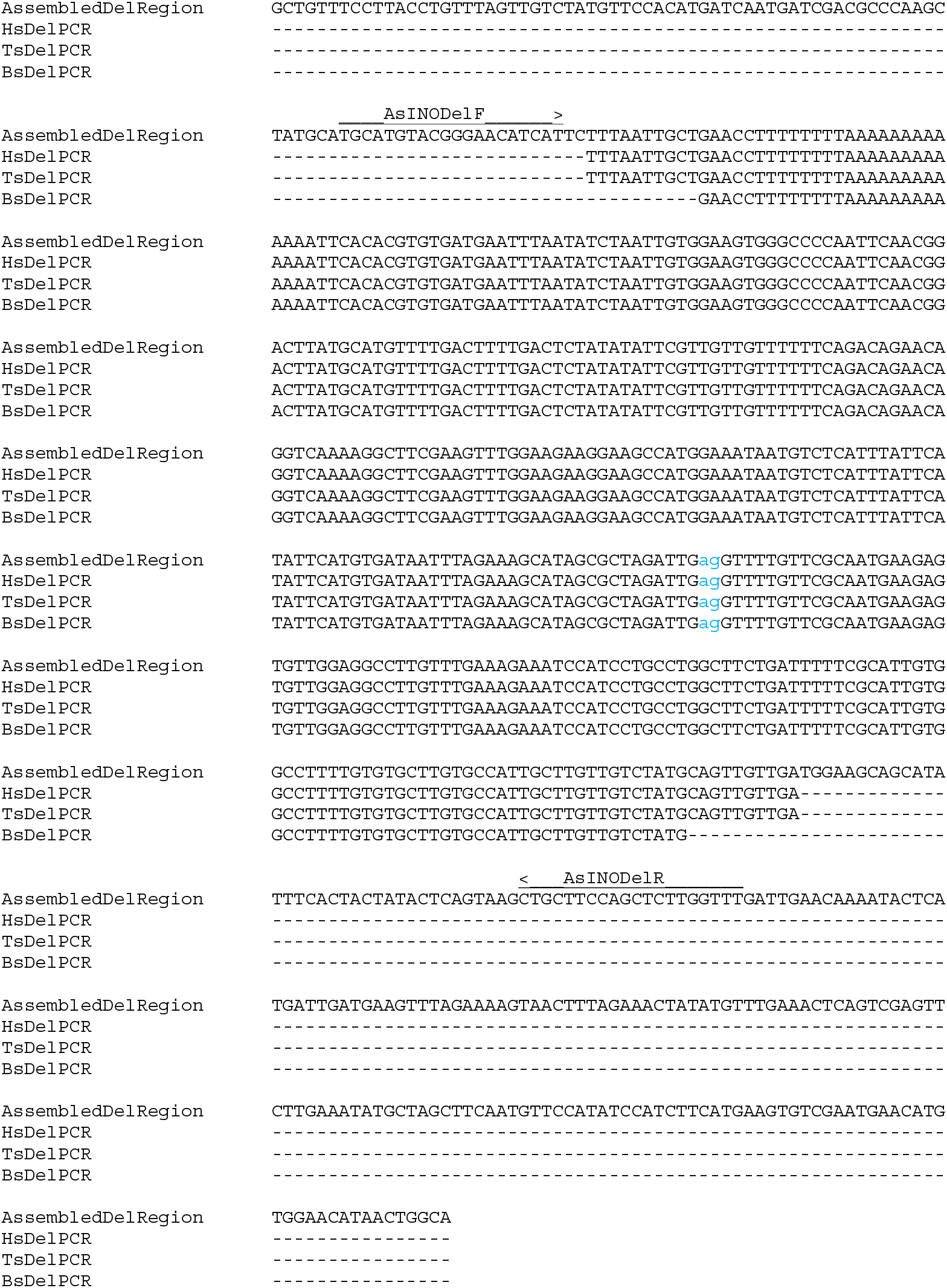
Aligned sequences of the deletion region junction. “AssembledDelRegion” is the sequence assembled from the Illumina reads selected from among all reads by BLAST searches with the reads flanking the deletion. The deletion falls between the “ag” nucleotides shown in lowercase light blue. The other sequences are those of PCR products produced using the AsINODel primers (indicated by arrows above the sequence) on DNA from Hs, Ts, or Bs as indicated, with sequence determined using the AsINODelR primer. All sequences show an identical deletion.

Primers were designed that would amplify the region spanning the deletion in mutant plant DNA (Figure 3). These primers (AsINODel F and AsINODel R) were used for PCR on DNA from Hs and Ts lines. Both lines produced a fragment with migration consistent with a length consistent with the 456 bp expected for the deletion (Figure 4A). Further, because the region between the primers in wild type is too large to be amplified in standard procedures, this procedure provides a positive assay for the presence of the deletion. In combination with primers that will amplify a region of the wild-type *INO* gene, this would enable a codominant test for the wild-type and seedless alleles of the gene in a single reaction. To test this, we modeled amplification from a “heterozygote” by mixing DNA from Hs and wild-type *A. squamosa*. We found that the amplification with the deletion-specific primers and LMINO1/2 primers in a single reaction could clearly differentiate the homozygous seedless, homozygous wild-type and heterozygous lines (Figure 4B).

**Figure 4.**
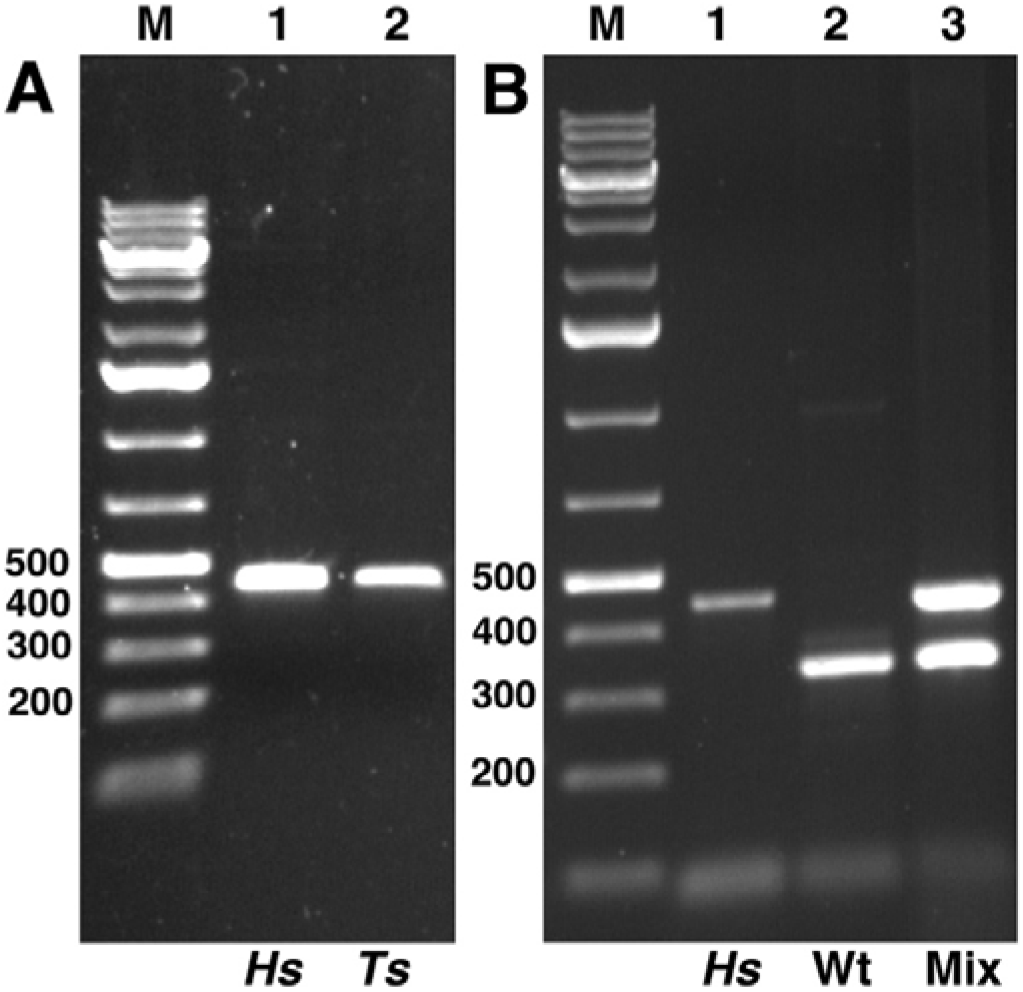
PCR of *INO* deletion region. A) AsINODel primers designed to amplify a fragment spanning the deletion junction were used on Hs and Ts DNA and the products were electrophoresed on an agarose gel. The Hs DNA produced the expected 457 bp fragment, and a comigrating fragment was produced from Ts. B) A combination of the AsINODel and LMINO1/2 primer pairs was used in single reactions to amplify DNA from Ts, wild-type and a mixture of wild-type and Ts (“Mix”). Only the expected 457 and 350 bp fragments were amplified from the Ts and wild-type DNA, respectively. Both bands were amplified from the mixture demonstrating effective detection of a heterozygous state.

Subsequent work showed that AsINODel primer PCR on all three seedless isolates produced comigrating fragments (Figure 4A and Figure 5A). The sequences of the PCR products were determined for all three seedless lines and were aligned with the assembled deletion region sequence (Figure 3). The sequence of the deletion junction was confirmed in the sequence from Hs, and an identical deletion was present in the same position in the sequences from mutants Bs and Ts (Figure 3), consistent with all lines producing PCR products of the same size. Further, the presence of identical deletions in all three lines indicates a single origin for the *INO* deletion among the isolates.

**Figure 5.**
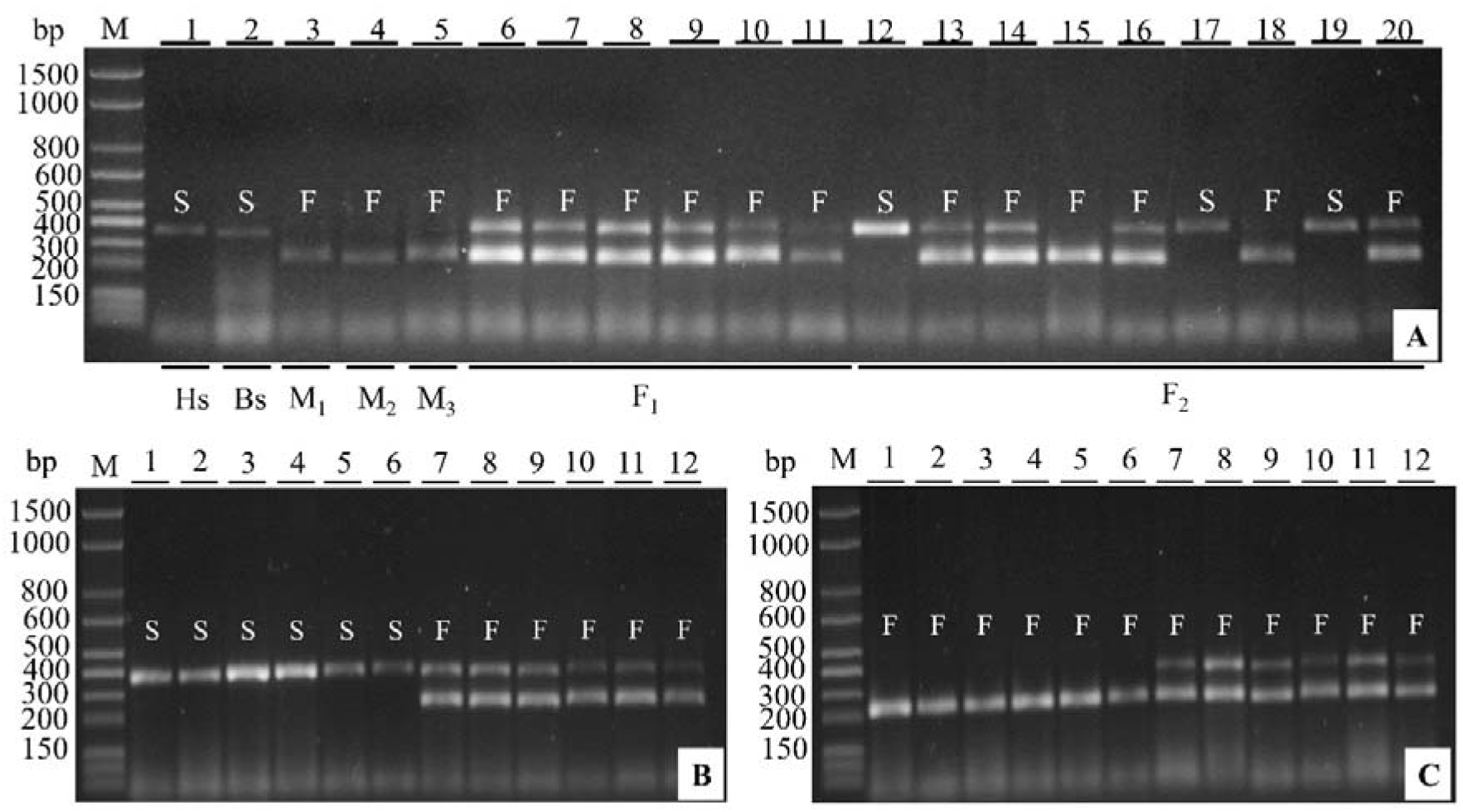
Amplifications products for combined AsINODel and LMINO1/2 primer pairs obtained from DNA samples of populations indicated below the agarose gel lanes (A). M bp: molecular weight marker in base pairs; Hs: Hawaiian seedless; Bs: Brazilian seedless; M_1_, M_2_, M_3_: wild-types; F_1_: first filial generation (lanes 6 to 11); F_2_: self-fertilization (lanes 12 to 20). Generations BC_Bs_ (B) and BC_M_ (C) lanes 1 to 12 contain the amplified fragments from the 12 individuals each. S: seedless; F: fertile (phenotypic determinations of the source plants).

### Codominant marker analysis

To evaluate use of the AsINODel primers together with the LMINO1/2 pair as codominant markers for plant breeding, lines M_1_, M_2_, and M_3_ were used as wild-type for *INO* gene and the presence of seeds, and the mutant Bs for the *INO* deletion and the absence of seeds. The genotype Hs was used as a positive control for the marker AsINODel. After PCR reactions, the combined primer pairs amplified a single fragment of 350 bp from the wild-type parents, representing the presence of the *INO* gene, and a 456 bp fragment from the mutants Bs and Hs, corresponding to the deletion junction fragment (Figure 5A).

As expected, the wild-type fertile parents (M_1_, M_2_ and M_3_) were homozygous for the presence, and the mutant seedless parent Bs was homozygous for the deletion of the *INO* gene. In generation F_1_ both *INO* gene and deletion region bands were present in all 32 genotypes evaluated, consistent with the expected heterozygous state (Table 3 and Figure 5A).

**Table 3.** Molecular analysis of codominant segregation of parents (M_1_, M_2_, M_3_ and Bs), F_1_ generations and generation F_2_, BC_M_ and BC_Bs_ backcrosses of the M_2_ family in *A*. *squamosa,* Janaúba-MG, Brazil.

| Parents and Generations | Observed proportion | | | Expected ratio | $\chi^2$ | p-value |
| --- | --- | --- | --- | --- | --- | --- |
|  | *INO INO* | *INO ino* | *ino ino* |  |  |  |
| Parents |  |  |  |  |  |  |
| M<sub>1</sub> | 1 |  |  |  |  |  |
| M<sub>2</sub> | 1 |  |  |  |  |  |
| M<sub>3</sub> | 1 |  |  |  |  |  |
| Bs |  |  | 1 |  |  |  |
| Generation F<sub>1</sub> |  |  |  |  |  |  |
| M<sub>1</sub> × Bs | 0 | 9 | 0 | 0:1:0 |  |  |
| M<sub>2</sub> × Bs | 0 | 17 | 0 | 0:1:0 |  |  |
| M<sub>3</sub> × Bs | 0 | 6 | 0 | 0:1:0 |  |  |
| Generation F<sub>2</sub> |  |  |  |  |  |  |
| (M<sub>2</sub> × Bs) × (M<sub>2</sub> × Bs) | 34 | 70 | 41 | 1:2:1 | 0.848 | 0.654<sup>ns</sup> |
| Backcrosses |  |  |  |  |  |  |
| BC<sub>M</sub> |  |  |  |  |  |  |
| F<sub>1</sub>(M<sub>2</sub> × Bs) × M<sub>2</sub> | 30 | 26 | 0 | 1:1 | 0.286 | 0.593<sup>ns</sup> |
| BC<sub>Bs</sub> |  |  |  |  |  |  |
| F<sub>1</sub>(M<sub>2</sub> × Bs) × Bs | 0 | 30 | 31 | 1:1 | 0.016 | 0.898<sup>ns</sup> |

<sup>ns</sup> Non-significant values at the 5% significance level, $\chi^2$ Chi-square test estimate.

The codominant markers were also used in genotyping of individuals of the F_2_ and backcross generations (BC_M_ and BC_Bs_ from M_2_×Bs) (Table 3) with some individuals selected to illustrate the discriminatory capacity of markers in Figure 5 A, B and C. In generation F_2_, analysis of 145 plants gave results consistent with the expected ratio of 1:2:1 for homozygous *INO*: heterozygous: homozygous *ino* deletion (Table 3). The expected ratios of genotypes were also observed for the sixty-one BC_Bs_ plants and the fifty-six BC_M_ plants, where ratios close to 1:1:0 and 0:1:1 were observed respectively (Figure 5B, 5C and Table 3). The χ^2^ test on each of these generations showed probabilities consistent with a monogenic segregation of the wild-type and mutant alleles.

Fruit was obtained from sixty-three of the genotyped segregating plants. In every case where a band corresponding to the wild-type *INO* gene was present (11 homozygous *INO* and 37 heterozygous plants) the seeded phenotype was observed. Similarly, all of the fifteen plants homozygous for the deletion allele exhibited the Bs mutant seedless phenotype. Thus, there was 100% correlation of the seeded and seedless phenotypes with the wild-type containing and homozygous *ino* deletion genotypes, respectively. To obtain more information to further characterize the linkage between the molecular and visible phenotypes we examined the ovules of plants that were flowering but not yet producing fruit where the *ino* mutant ovules could be differentiated from wild-type (Supplemental Figure S2). This determination was done on ninety-five plants and once again there was a complete correspondence between the molecular genotype and the presence of wild-type ovules or aberrant ovules incapable of forming seeds (Supplemental data file 1).

If the *ino* mutant gene is only linked to the seedless mutant allele, and not causally responsible for it, then recombination between the molecular and phenotypic markers would be possible. Among segregating F_2_ plants, a single recombination event could be observed in the wild-type *INO* containing chromosome of heterozygous progeny, or in either chromosome of homozygous *ino* progeny. The same would be true of chromosomes deriving from the F_1_ plants in the BC Bs progeny. In these cases, recombination would lead to a switch of the seedless/seeded phenotype and breakage of the 100% cosegregation (Supplemental Methods). Seed or ovule phenotypic data were produced for 57 informative heterozygous plants and 36 homozygous mutant plants (allowing examination of a total of 114 chromosomes) (Supplemental data file 1). Thus, no recombination events were observed between the *INO* locus and the Bs gene in 114 chromosomes. This allows an initial limit on the maximum genetic distance between *ino* and Bs. A genetic distance of 3.5 cM would lead to a prediction of 3.5% recombination, where 0% recombination was observed. The comparison of these observed and expected result using the χ^2^ test produces χ^2^ = 3.99 and a corresponding P-value of 0.046. Thus, a genetic distance of 3 cM (or greater) can be rejected at the 5% probability level (Supplemental data file 1). In addition, within the 587 kb of sequence surrounding the *INO* gene, we find that the 16 kb deletion is the only significant difference between wild-type and Hs in this region (the only other being a single base change in an intron of a predicted gene). Together these observations are consistent with the *INO* gene deletion being the lesion responsible for the seedless phenotype.

### Genetic diversity using SSR markers

Of the 67 SSR primers used, 41 amplified under the tested conditions, 12 presented a null amplification pattern with no bands evident, and the remaining 14 did not present a clear band pattern in the tested samples (Supplemental Table S4). Among the primers that amplified, 38 generated a total of 63 monomorphic alleles; only three primers (LMCH 3, 39, and 137) generated specific polymorphisms among the genotypes evaluated.

The LMCH 3 primer generated two alleles between 200 and 300 bp for the accession M_2_ genotype and the seedless genotype, respectively. The primer LMCH 39 amplified five alleles in total. Three generated a band only in genotype M_2_, distinguishing it from seedless genotypes. Similarly, the primer LMCH 137 amplified five alleles. However, in this marker the seedless genotypes presented two specific bands differing from genotype M_2_, displaying amplification of two monomorphic bands common to all individuals. The genetic distance between genotypes, estimated by the complement of the similarity matrix generated by the Jaccard index, ranged from 0 among genotypes Bs, Ts, and Hs, and 0.0933 between the M_2_ with Bs, Ts, and Hs. Thus, these markers differentiated between the mutant lines and the wild-type line, but could not demonstrate any divergence between the three mutant accessions.

## Discussion

Three seedless stenospermocarpous sugar apple varieties, Bs, Ts, and Hs, from differing global locales have been described [Lora et al. (2011), Santos et al. (2014) and this work]. The seedless phenotype results from an ovule defect (Lora et al., 2011, Santos et al., 2014) indistinguishable from the ovule effects of the *inner no outer* mutation in *Arabidopsis* (Villanueva et al., 1999), and was associated with a deletion of this locus in the Ts accession (Lora et al., 2011). On this basis, Lora et al. (2011) hypothesized that the deletion was the cause of the seedless trait, but segregation data were not available. We show that the three seedless accessions all carry identical 16 kb deletions of the region including *INO*. The identical sequence of the deletion region in the three accessions indicates that the mutant allele arose as a single deletion event in a common ancestor and propagated globally. The deletion was not associated with any apparent repeated sequence at the deletion junction, but was a precise excision of the deleted region. We tested the causal relationship between the deletion and seedlessness by examining segregation of the seedless trait and the deletion. We demonstrated that seedlessness is a single locus recessive trait and that the fertile/seedless phenotypes cosegregated 100% with the presence/absence of the *INO* gene among progeny plants (114 evaluated chromosomes). Our data demonstrate that the *ino* deletion and the seedless trait are separated by less than 3.5 cM. We have determined the sequence of 587 kb surrounding the deletion region, and based on prior measures of the relationship between genetic and physical mapping distances (Puchta and Hohn, 1996) this would constitute from three to thirty centimorgans and so would include the seedless mutation. Outside of the deletion that includes the *INO* gene, we found no other significant differences from wild type in the sequenced region surrounding the *INO* locus. Together with the clear expectation of an *ino* gene deletion causing the observed ovule phenotype, these data support the *ino* deletion being the sole cause of the seedless trait.

It is possible that the deletion and seedless trait was retained in multiple lines by selection despite significant outcrossing, or alternatively that all lines with this deletion are vegetatively propagated clones. Genotyping through the use of SSR markers failed to detect polymorphism among our available seedless accessions (Bs, Hs, Ts) but did show a low level of polymorphism (4.7%) when compared to the wild-type M_2_ (indicating low genetic diversity among genotypes). These data therefore do not rule out a possible spread by vegetative propagation. The utilized SSR markers were developed for *A. cherimolia* and no specific SSR primers were available for *A. squamosa* and some outbreeding could have been missed in this analysis. But with the current data the simplest explanation is that a single mutational event was dispersed to different continents through vegetative propagation. Tracing a temporal and geographical route of this dispersal is difficult since the species has been widely developed and naturalized.

It is believed that the primary center of origin of *A. squamosa* is in the lowlands of Central America. Historical data indicate accessions to Mexico by the natives of the region (Pinto et al., 2005). According to these authors, after their arrival, the Spaniards were responsible for spreading the seeds to the Philippines. By 1590, the species had been introduced into India. In Brazil, 1626 is the first record of introduction of the species. By the early 17th century the species was already widespread in Indonesia, China, Australia, Polynesia and Hawaii. In 1955, the Cuban seedless variety had been introduced in the state of Florida in the United States. The mutant Brazilian accession Bs was first described in 1940 in the state of São Paulo (Cunha, 1953). We obtained Hs from a commercial nursery in Hawaii. The nursery reports obtaining the line from the Philippines. Their source in the Philippines reports that the line was brought to the Philippines from an unknown location in the 1920s, but documentary evidence of this was not found. Information on the origin of Ts was not available.

Lora et al. (2011) designed primers within, or closely flanking the transcribed region of *INO* for detection of the wild-type locus, and demonstrated amplification from wild-type *A. squamosa, A. cherimola,* and four other species that cover the phylogenetic range of the genus *Annona*. These primers were confirmed to function as expected in our *A. squamosa* varieties (Nassau et al., 2021 and this work). Notably the primer pairs only allow detection of the wild-type locus and a homozygous mutant could only be detected by the absence of amplification. According to Singh and Singh (2015), markers that can directly identify the traits of interests should be more efficient in incorporating and monitoring genes in breeding programs. Our determination of the sequence of the region from which *INO* has been deleted enabled formulation of primers (AsINODel) for direct detection of the mutant allele allowing codominant detection of both alleles in wild-type and seedless mutant parents and to differentiate fertile heterozygotes from homozygous fertile plants in a single reaction.

The improvement of perennial plants, especially fruit species, is a challenging task due to a long juvenile period and seed dormancy, among other factors, which translates into a relatively long generation period (McClure et al., 2014). In *A. squamosa,* for example, the beginning of flowering of adult plants, originated from seeds, under specific cultivation conditions, would take at least three years (Pereira and Borém, 2021) until the characterization of the fruits, limiting the number of genotypes that can be studied, which makes the breeding process slow and expensive. The PCR assay using a combination of the primers detecting the wild-type and mutant alleles allows plants to be genotyped in the seedling stage, accelerating the breeding program. Given the generality of the *INO* gene detecting primers for use in multiple species of *Annona* (Lora et al., 2011), and the fertility of interspecific crosses in this genus, our codominant genotyping is applicable to introgression of the seedless trait into elite sugar apple varieties and into other cultivated *Annona* species.

## Conclusions

The presence/absence of seeds in *A. squamosa* displays a monogenic inheritance. The *INO* allele, whether homozygous or heterozygosis (*INO*_—_), conditions the presence of seeds and has complete dominance over the recessive seedless condition (*ino ino*). The Bs, Ts and Hs mutant accessions all display the same deletion sequence for the *INO* locus, demonstrating a common origin of genotypes. The deletion is simple, not resulting in insertion or duplication of any sequence at the deletion junction. Determination of the deletion sequence allowed formulation of primers (AsINODel) that specifically detect the deletion. The AsINODel primers together with the marker LMINO1/2 display co-dominance and can distinguish dominant homozygotes from heterozygotes. The validated LMINO and AsINODel primers can facilitate introgression of the seedless trait in *A. squamosa* and other members of this genus.

## Acknowledgments

We thank Trang Dang for technical support and Debra Skinner for help with planning and analysis for sequence determination.

## Author Contributions

Experiments were conceived by S.N., S.P and C.S.G. Crossing and genetic analyses were performed by B.R.A.R. with help from M.C.T.P. Whole genome sequence analysis was performed by C.S.G. and B.R.A.R. S.N., B.R.A.R and C.S.G. wrote the manuscript. The author responsible for distribution of materials integral to the findings presented in this article in accordance with the policy described in the Instructions for Authors (https://academic.oup.com/plphys/pages/general-instructions) is: Silvia Nietsche

---

**Supplemental Figure S1.**
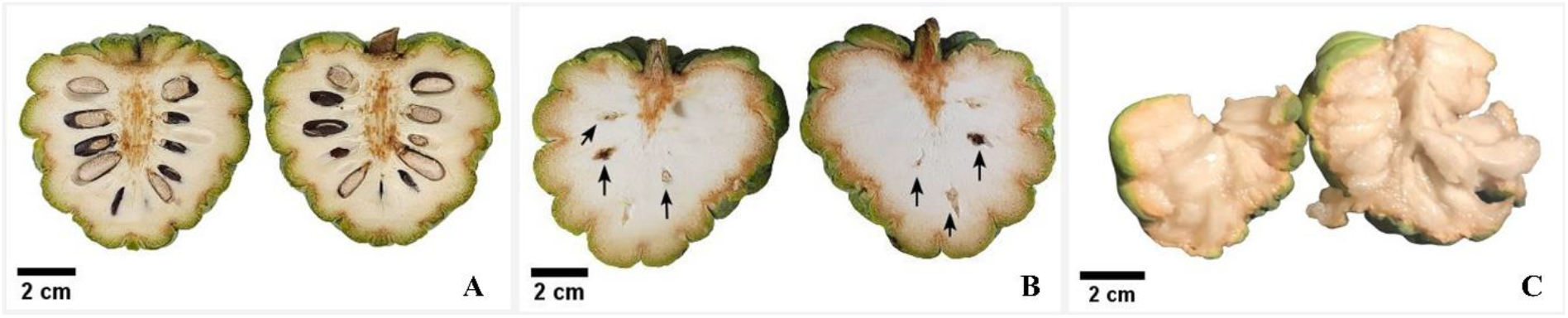
Fruit of *A. squamosa.* Longitudinal sections of fruit from (A) M_2_ wild type and (B) the mutant Brazilian seedless (Bs) showing seed rudiments (arrows), and (C) split fruit from Thai seedless (Ts). Backgrounds of images were normalized to white for clarity.

**Supplemental Figure S2.**
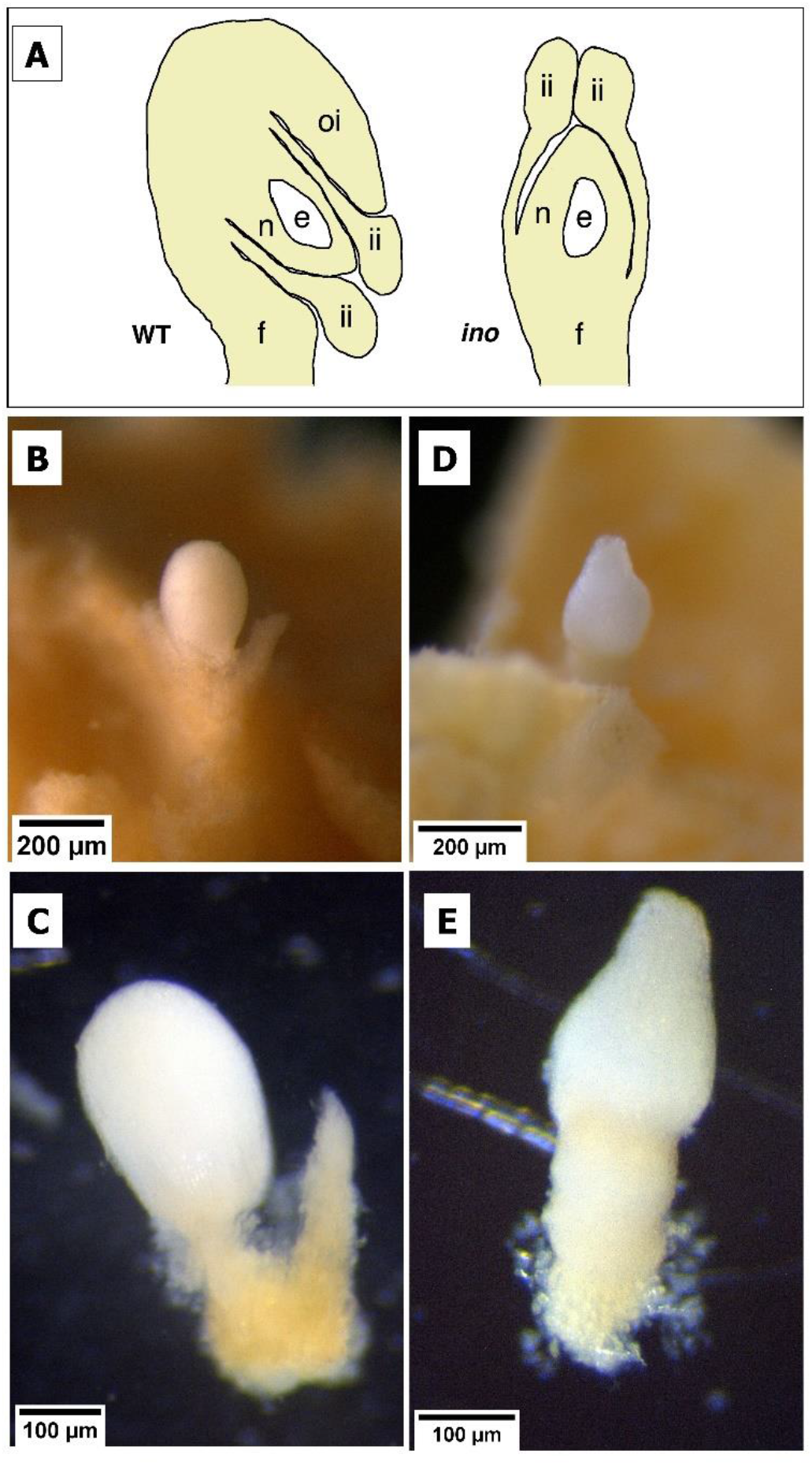
Determination of ovule phenotype on freshly dissected flowers. (A) Diagrams of *A. squamosa* ovule structure with sections of wild-type (left) and ino mutant (right) ovules redrawn based on images in Lora et al. (2011). (B to E) Images of freshly dissected *A. squamosa* ovules photographed with a stereomicroscope (Leica, M205C) using the Leica Application Suite software (LAS v4.11). (B and C) M_2_ wild type. (D and E) mutant Brazilian seedless. The domed shape of the wild-type and the more pointed-erect shape of the *ino* mutant are easily differentiated. e, embryo sac; f, funiculus; ii, inner integument; n, nucellus; oi, outer integument.

**Supplemental Figure S3.**
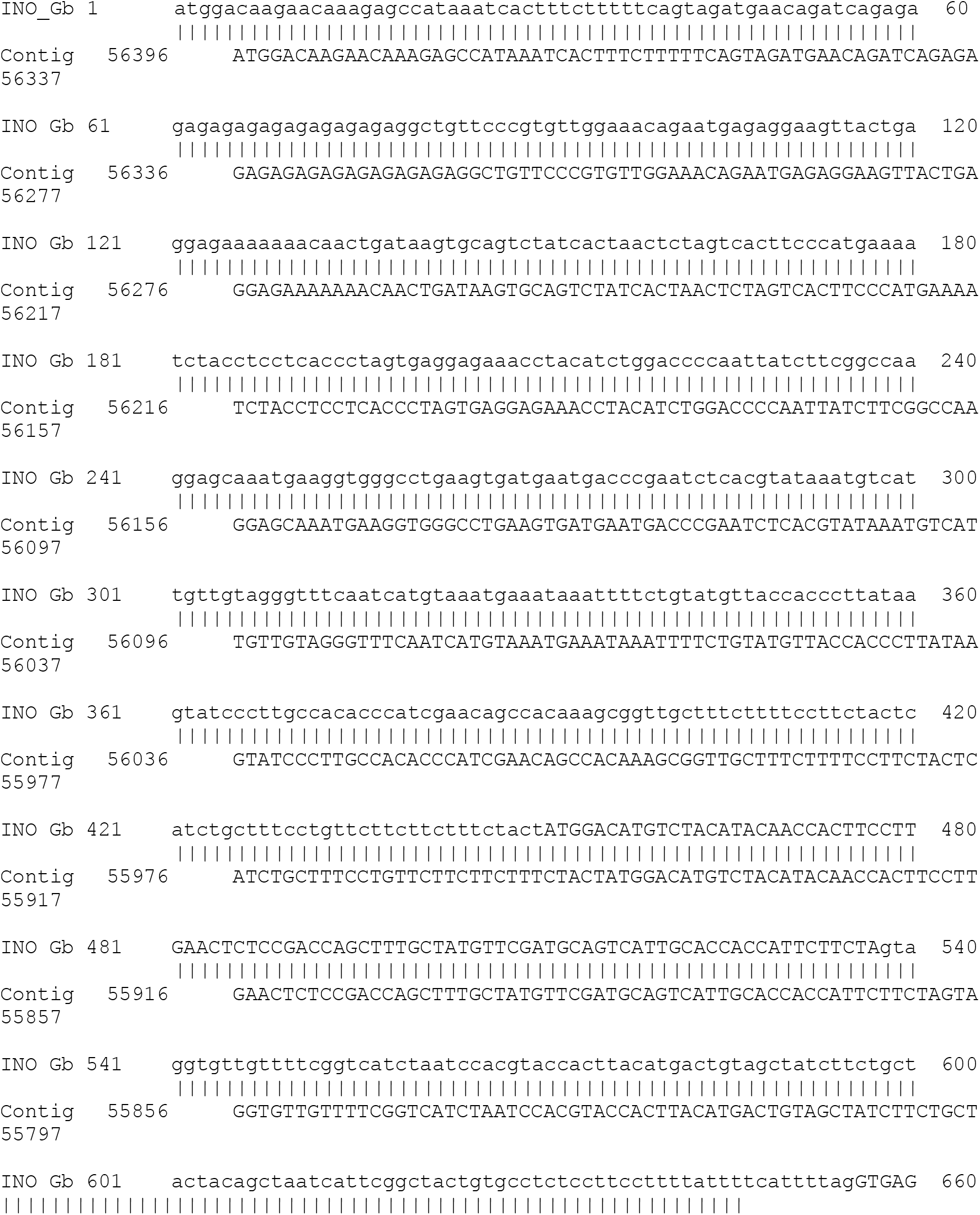
Alignment produced by BLAST search of whole genome assembly with the previously determined *A. squamosa INO* gene sequence (GenBank GU828033.1). “INO Gb” is the sequence from GenBank with noncoding flanking sequences and introns in lower case and coding regions in upper case. “Contig” is sequence from the single 587 kb contig (#00001115) identified by BLAST as containing the aligning sequence. The Contig sequence differs from the GenBank entry only in the presence of a 40 bp insertion in the fourth intron, and two single base insertions in the last dozen residues of the alignment over the entire 1847 bp aligned sequence.

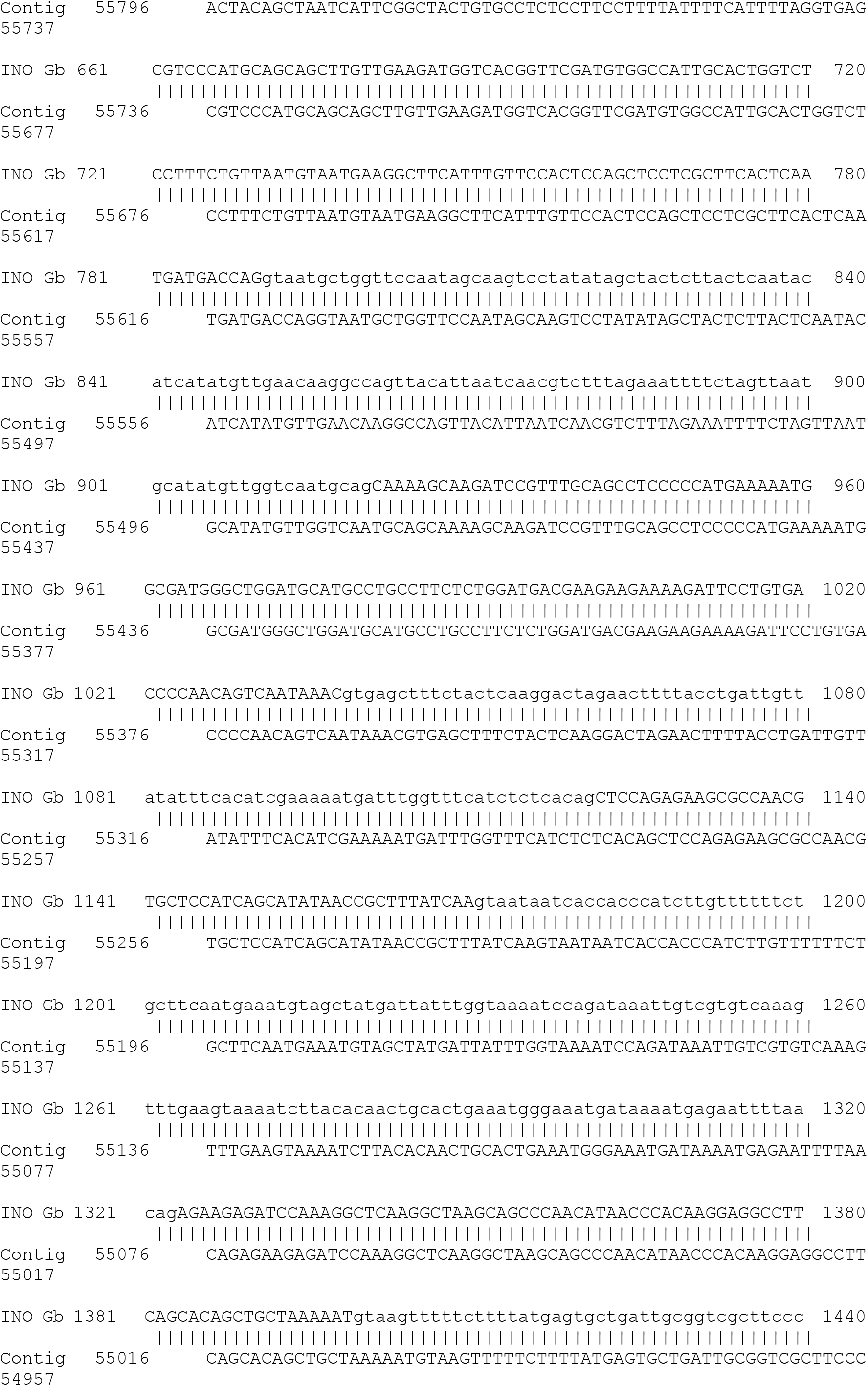

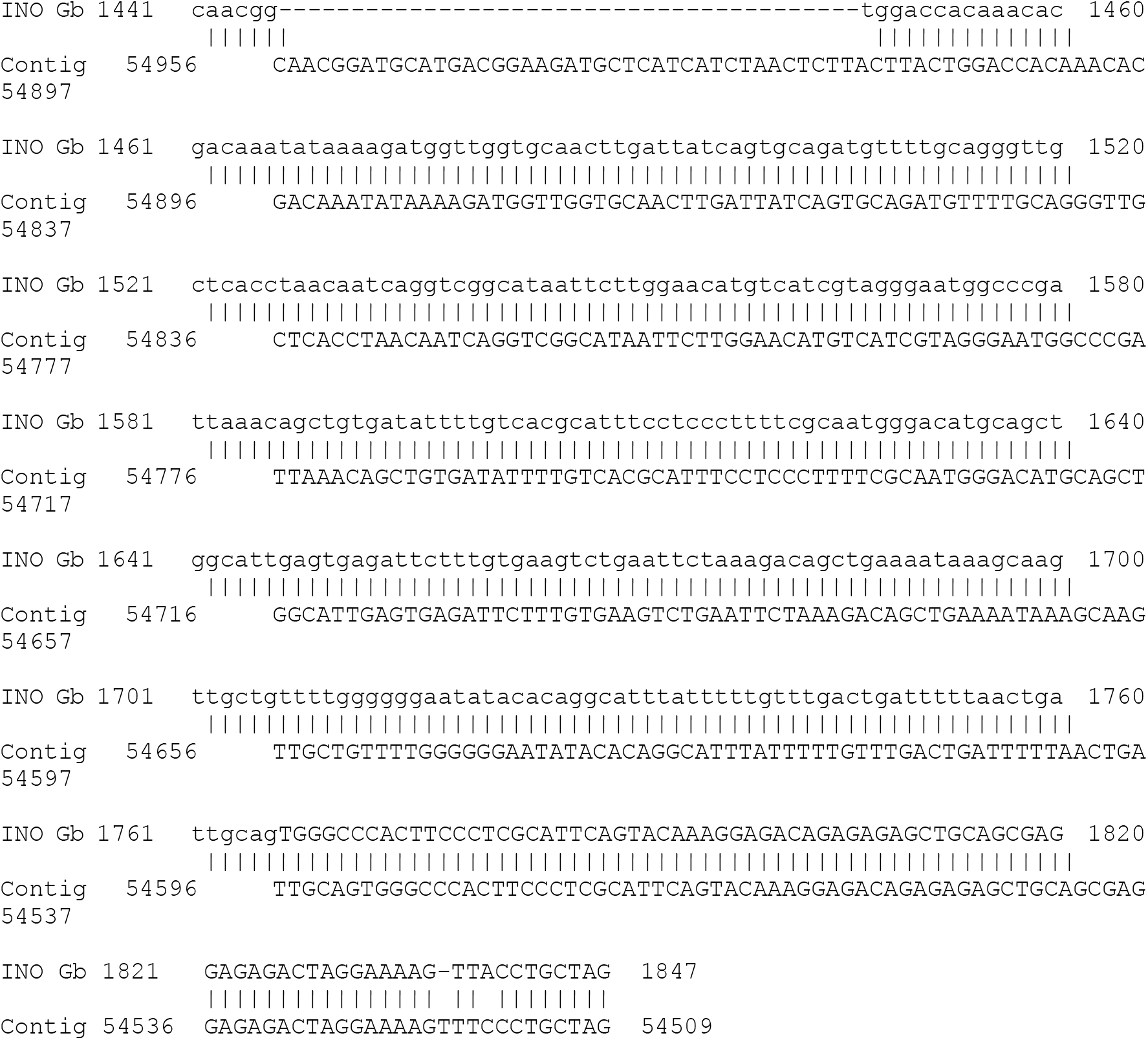

**Supplemental Figure S4.**
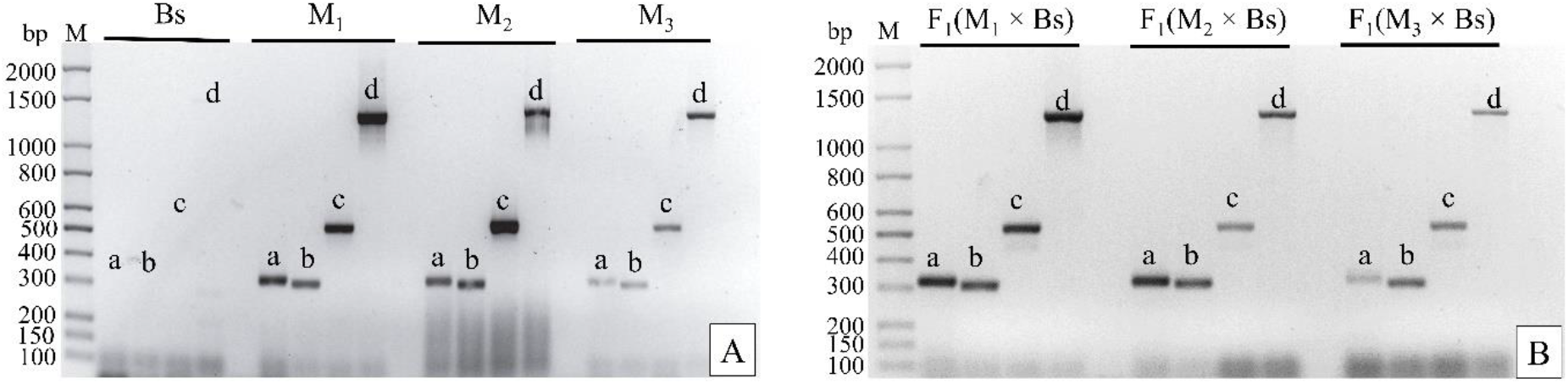
Amplification products for primer pairs LMINO. LMINO 1/2 (a), LMINO3/4 (b), LMINO5/6 (c), and LMINO7/8 (d) obtained from DNA samples from each of the parents (A) and of the three F_1_ progenies selected for conducting the segregating generations (B), indicated below the 1.2% agarose gels in TBE 1× buffer. M bp: molecular weight markers in base pairs; Bs: Brazilian seedless; M_1_, M_2_, and M_3_: wild-types (fertiles).

