## Supplemental tables for "Seedless fruit in *Annona squamosa* L. is monogenic and conferred by *INO* locus deletion in multiple accessions"

**Supplemental Material: Tables 1 - 4**

**Supplemental Table S1.** Oligonucleotides used in the genetic inheritance studies.

Identification of primers, sequence and product size in base pairs (bp).

| **Primer** | **Primer sequence (5' - 3')** | **bp** |
| --- | --- | --- |
| LMINO1  LMINO2 | CCTAAATGAAGGGTTTACATGTGGC  GCCCACCTTCATTTGCTCCTTGG | 350 |
| LMINO3  LMINO4 | ATGGACATGTCTACATACAACCAC  CGAGGAACTGCAGTGGAACAAATG | 350 |
| LMINO5  LMINO6 | AGAAGAGATCCAAAGGCTCAAGG  GCAGCTCTCTCTATCTCCTTTG | 600 |
| LMINO7  LMINO8 | GCGTGTAAGCAGATTCCCTTTACC  AGCGTCCCATGCAGCAGCTTGTT | 1200 |
| AsINODelF  AsINODelR | AAACCAAGAGCTGGAAGCAG  TGCATGTACGGGAACATCAT | 456 |

LMINO primers from Lora et al. (2011), AsINODel primers, this work.

**Supplemental Table S2.** Phenotypic segregation hypotheses.

Expected in generations F_2_ for one, two, and three genes involved in seed formation in *Annona squamosa* obtain in Janaúba-MG, Brazil, in 2020.

| **Generation F_2_** | **Hypothesis** |  | **Observed proportion** | |  | **χ²** | **P (%)** |
| --- | --- | --- | --- | --- | --- | --- | --- |
|  |  |  | **Presence** | **Absence** |  |  |  |
| M_2_ | 3:1 |  | 190 | 71 |  | 2.039 | 15.325^ns^ |
| M_2_ | 9:7 |  | 190 | 71 |  | 17.220 | 0.003 |
| M_2_ | 13:3 |  | 190 | 71 |  | 0.125 | 72.304^ns^ |
| M_2_ | 15:1 |  | 190 | 71 |  | 11.526 | 0.068 |
| M_2_ | 37:27 |  | 190 | 71 |  | 15.408 | 0.008 |
| M_2_ | 63:1 |  | 190 | 71 |  | 90.815 | 0.000 |

^ns^ Non-significant values at the 5% significance level, χ² Chi-square test estimate.

**Supplemental Table S3.** Genotypic segregation hypotheses.

Expected in generations F_2_ for one, two, and three genes involved in seed formation in *Annona squamosa* obtain in Janaúba-MG, Brazil, in 2020.

| **Generation F_2_** | **Hypothesis** |  | **Observed proportion** | |  | **χ²** | ***p*-value** |
| --- | --- | --- | --- | --- | --- | --- | --- |
|  |  |  | **Presence** | **Absence** |  |  |  |
| M_1_ | 3:1 |  | 99 | 43 |  | 2.11 | 0.1461^ns^ |
| M_1_ | 9:7 |  | 99 | 43 |  | 10.47 | 0.0012 |
| M_1_ | 13:3 |  | 99 | 43 |  | 12.40 | 0.0004 |
| M_1_ | 15:1 |  | 99 | 43 |  | 139.96 | 0.00 |
| M_1_ | 37:27 |  | 99 | 43 |  | 8.25 | 0.0041 |
| M_1_ | 63:1 |  | 99 | 43 |  | 761.47 | 0.00 |
| M_2_ | 3:1 |  | 190 | 71 |  | 0.68 | 0.4111^ns^ |
| M_2_ | 9:7 |  | 190 | 71 |  | 29.04 | 0.00 |
| M_2_ | 13:3 |  | 190 | 71 |  | 12.24 | 0.0005 |
| M_2_ | 15:1 |  | 190 | 71 |  | 195.56 | 0.00 |
| M_2_ | 37:27 |  | 190 | 71 |  | 24.03 | 0.00 |
| M_2_ | 63:1 |  | 190 | 71 |  | 1115.62 | 0.00 |
| M_3_ | 3:1 |  | 69 | 33 |  | 2.94 | 0.0864^ns^ |
| M_3_ | 9:7 |  | 69 | 33 |  | 5.38 | 0.0203 |
| M_3_ | 13:3 |  | 69 | 33 |  | 12.39 | 0.0004 |
| M_3_ | 15:1 |  | 69 | 33 |  | 118.61 | 0.00 |
| M_3_ | 37:27 |  | 69 | 33 |  | 4.05 | 0.0443 |
| M_3_ | 63:1 |  | 69 | 33 |  | 628.71 | 0.00 |

^ns^ Non-significant values at the 5% significance level, χ² Chi-square test estimate.

**Supplemental Table S4.** Oligonucleotides used in the diversity study.

Description of primers, with their amplification repeats, annealing temperature (Ta), number of amplified alleles (A), number of polymorphic alleles (P) in genotypes of *A. squamosa* with and without seeds obtained in Janaúba-MG, Brazil, 2020.

| **SSR** | **Repetition** | **Ta°C** | **A** | **P** | **Source** |
| --- | --- | --- | --- | --- | --- |
| LMCH 1 | (CT)20 | 55 | 1 | 0 | Escribano et al., 2004 |
| LMCH 2 | (CCT)5 | 55 | 1 | 0 | Escribano et al., 2004 |
| LMCH 3 | (GA)13 | 55 | 2 | 2 | Escribano et al., 2004 |
| LMCH 4 | (GA)14 | 55 | 1 | 0 | Escribano et al., 2004 |
| LMCH 5 | (CT)10 | 55 | - | - | Escribano et al., 2004 |
| LMCH 6 | (CT)14 | 55 | - | - | Escribano et al., 2004 |
| LMCH 7 | (GA)9 | 55 | - | - | Escribano et al., 2004 |
| LMCH 8 | (GA)8 | 55 | 0 | 0 | Escribano et al., 2004 |
| LMCH 9 | (GA)6 | 55 | 1 | 0 | Escribano et al., 2004 |
| LMCH 10 | (CT)12 | 55 | - | - | Escribano et al., 2004 |
| LMCH 11 | (CT)10 | 55 | - | - | Escribano et al., 2004 |
| LMCH 12 | (CT)17 | 55 | 5 | 0 | Escribano et al., 2004 |
| LMCH 13 | (CT)39 | 55 | - | - | Escribano et al., 2004 |
| LMCH 14 | (GA)8 | 55 | 7 | 0 | Escribano et al., 2004 |
| LMCH 16 | (GA)20 | 55 | - | - | Escribano et al., 2004 |
| LMCH 29 | (GA)9 | 45 | 1 | 0 | Escribano et al., 2008 |
| LMCH 33 | (CT)11 | 50 | 2 | 0 | Escribano et al., 2008 |
| LMCH 34 | (GA)11 | 50 | 0 | 0 | Escribano et al., 2008 |
| LMCH 36 | (GA)10 | 50 | 1 | 0 | Escribano et al., 2008 |
| LMCH 37 | (GA)15 | 50 | 0 | 0 | Escribano et al., 2008 |
| LMCH 38 | (GA)22 | 55 | - | - | Escribano et al., 2008 |
| LMCH 39 | (CT)11 | 55 | 5 | 3 | Escribano et al., 2008 |
| LMCH 40 | (GA)17 | 55 | 0 | 0 | Escribano et al., 2008 |
| LMCH 42 | (GA)11 | 55 | 1 | 0 | Escribano et al., 2008 |
| LMCH 43 | (GA)9 | 55 | 1 | 0 | Escribano et al., 2008 |
| LMCH 48 | (GA)12 | 55 | 2 | 0 | Escribano et al., 2008 |
| LMCH 53 | (GA)8 | 55 | 1 | 0 | Escribano et al., 2008 |
| LMCH 54 | (GAA)6 | 50 | 2 | 0 | Escribano et al., 2008 |
| LMCH 57 | (CT)5 | 55 | 4 | 0 | Escribano et al., 2008 |
| LMCH 63 | (GA)13 | 50 | 4 | 0 | Escribano et al., 2008 |
| LMCH 68 | (GA)30 | 55 | 2 | 0 | Escribano et al., 2008 |
| LMCH 69 | (GA)9/(GT)3 | 55 | 1 | 0 | Escribano et al., 2008 |
| LMCH 70 | (GA)8 | 55 | 1 | 0 | Escribano et al., 2008 |
| LMCH 71 | (GA)14 | 55 | 1 | 0 | Escribano et al., 2008 |
| LMCH 72 | (GA)8 | 55 | 1 | 0 | Escribano et al., 2008 |
| LMCH 73 | (CT)13 | 55 | 1 | 0 | Escribano et al., 2008 |
| LMCH 78 | (GA)9 | 55 | 1 | 0 | Escribano et al., 2008 |
| LMCH 79 | (CT)12 | 55 | 1 | 0 | Escribano et al., 2008 |
| LMCH 80 | (GA)15 | 55 | 2 | 0 | Escribano et al., 2008 |
| LMCH 83 | (CT)36 | 55 | 1 | 0 | Escribano et al., 2008 |
| LMCH 87 | (GA)15 | 55 | 1 | 0 | Escribano et al., 2008 |
| LMCH 88 | (CT)17 | 55 | 0 | 0 | Escribano et al., 2008 |
| **SSR** | **Repetition** | **Ta °C** | **A** | **P** | **Source** |
| LMCH 89 | (CT)11 | 55 | 2 | 0 | Escribano et al., 2008 |
| LMCH 90 | (CT)11 | 55 | - | - | Escribano et al., 2008 |
| LMCH 91 | (CT)11 | 55 | 1 | 0 | Escribano et al., 2008 |
| LMCH 92 | (CT)8 | 55 | 2 | 0 | Escribano et al., 2008 |
| LMCH 93 | (GA)8 | 55 | 2 | 0 | Escribano et al., 2008 |
| LMCH 96 | (CT)10 | 55 | 0 | 0 | Escribano et al., 2008 |
| LMCH 98 | (GA)10 | 55 | 0 | 0 | Escribano et al., 2008 |
| LMCH 102 | (CT)13 | 55 | 1 | 0 | Escribano et al., 2008 |
| LMCH 103 | (GA)19 | 55 | 0 | 0 | Escribano et al., 2008 |
| LMCH 106 | (GA)13 | 55 | 0 | 0 | Escribano et al., 2008 |
| LMCH 108 | (GA)9 | 55 | 0 | 0 | Escribano et al., 2008 |
| LMCH 109 | (GA)7 | 55 | 1 | 0 | Escribano et al., 2008 |
| LMCH 112 | (GA)12 | 55 | - | - | Escribano et al., 2008 |
| LMCH 114 | (CA)4/(CT)9 | 55 | - | - | Escribano et al., 2008 |
| LMCH 115 | (GA)13 | 50 | - | - | Escribano et al., 2008 |
| LMCH 119 | (GA)12 | 55 | - | - | Escribano et al., 2008 |
| LMCH 122 | (GA)9 | 55 | 1 | 0 | Escribano et al., 2008 |
| LMCH 127 | (CT)9 | 55 | 1 | 0 | Escribano et al., 2008 |
| LMCH 128 | (GA)11 | 55 | 2 | 0 | Escribano et al., 2008 |
| LMCH 131 | (GA)10 | 55 | 0 | 0 | Escribano et al., 2008 |
| LMCH 134 | (CT)8 | 48 | 0 | 0 | Escribano et al., 2008 |
| LMCH 137 | (GA)24 | 48 | 5 | 2 | Escribano et al., 2008 |
| LMCH 139 | (CT)9 | 55 | 1 | 0 | Escribano et al., 2008 |
| LMCH 142 | (CT)12 | 55 | - | - | Escribano et al., 2008 |
| LMCH 144 | (CT)12 | 55 | 1 | 0 | Escribano et al., 2008 |

-Absence of band pattern in the samples tested.
